# Dengue activates mTORC2 signaling to counteract apoptosis and maximize viral replication

**DOI:** 10.1101/2020.09.22.308890

**Authors:** Christoph C. Carter, Jean Paul Olivier, Alexis Kaushansky, Fred D. Mast, John D. Aitchison

## Abstract

The mechanistic target of rapamycin (mTOR) functions in at least two distinct complexes: mTORC1, which regulates cellular anabolic-catabolic homeostasis, and mTORC2, which is an important regulator of cell survival and cytoskeletal maintenance. mTORC1 has been implicated in the pathogenesis of flaviviruses including dengue, where it contributes to the establishment of a pro-viral autophagic state. In contrast, the role of mTORC2 in viral pathogenesis is unknown. In this study, we explore the consequences of a physical protein-protein interaction between dengue non-structural protein 5 (NS5) and host cell mTOR proteins during infection. Using shRNA to differentially target mTORC1 and mTORC2 complexes, we show that mTORC2 is required for optimal dengue replication. Furthermore, we show that mTORC2 is activated during viral replication, and that mTORC2 counteracts virus-induced apoptosis, promoting the survival of infected cells. This work reveals a novel mechanism by which the dengue flavivirus can promote cell survival to maximize viral replication.

## INTRODUCTION

Mechanistic target of rapamycin (mTOR) is an essential serine/threonine kinase that functions in several key aspects of mammalian cell biology (reviewed in (1)). mTOR exerts its actions as a component of at least two distinct complexes, mTORC1 and mTORC2. mTORC1 is composed of five proteins: mTOR, Raptor, mLST8, PRAS40 and Deptor. mTORC1 functions as a master regulator of anabolic/catabolic homeostasis. In conditions of nutritional abundance, mTOR phosphorylates p70 ribosomal protein S6K (S6K) and eukaryotic initiation factor 4E binding protein 1 (4E-BP1), leading to increased protein synthesis. In conditions of nutrient scarcity mTORC1 is inactivated, stimulating autophagy, which allows for the recycling and turnover of cellular organelles and protein complexes (1).

mTORC2 is composed of six proteins: mTOR, Rictor, mLST8, SIN1, PRRL5L and Deptor. mTORC2 has distinct roles from mTORC1 in cellular physiology, but these roles are less well understood than those of mTORC1. mTORC2 promotes cell survival and proliferation through phosphorylation of AKT at ser473 (2). It is also known to play a role in the maintenance of the actin cytoskeleton, and when inactivated results in morphological abnormalities in some cell lines (3, 4). It has been suggested that mTORC2 may modulate translational machinery due to its association with ribosomes; however, the ramifications of this interaction are not well understood (5). Although mTORC1 is the canonical regulator of autophagy, mTORC2 has been implicated in the regulation of specific autophagic processes such as chaperone-mediated autophagy and mTORC1-independent autophagy (6, 7).

Given the important role of mTORC1 in regulating cellular metabolism, it is not surprising that several viruses have evolved mechanisms to modulate mTORC1 signaling (8, 9). Numerous viruses have been shown to manipulate mTORC1 activity during infection; some viruses activate mTORC1 to maintain cellular anabolic machinery, whereas others suppress mTORC1 activity to favor cap-independent viral protein synthesis (8, 9). In the case of dengue, virus-induced modulation of mTORC1 has been suspected due to the importance of autophagy in dengue infection (10–13), although this interaction of the virus with mTORC1 has not been comprehensively investigated. The importance of mTORC1 in dengue replication is supported by studies demonstrating increased viral replication in the response to pharmacologic mTOR inhibition and a recent study implicating mTORC1 in the dengue-induced activation of lipophagy (14, 15).

In contrast to mTORC1, the role of mTORC2 in virus-host interaction is poorly understood. Activation of mTORC2 has been documented in human cytomegalovirus, West Nile and influenza infection (16–18), but the functional role of mTORC2 in these infections remains unknown. Furthermore, to our knowledge no role for mTORC2 has been described in dengue infection.

Here, we describe a role for mTORC2 in promoting cell survival during dengue infection. We find that the dengue non-structural protein 5 (NS5) interacts with mTORC1 and mTORC2 complexes, and that dengue infection leads to the activation of mTORC2 signaling. We report that inactivation of mTORC2 signaling leads to a decrease in viral replication and an increase in virus-induced apoptosis and cell death. These findings suggest a mechanism by which dengue counteracts apoptosis to maintain cell survival and maximize viral replication.

## RESULTS

### Dengue NS5 protein interacts with mTORC1 and mTORC2

A previous quantitative proteomics approach (I-DIRT; Isotopic Differentiation of Interactions as Random or Targeted(19)), designed to identify *bona fide* dengue-host protein-protein interactions, defined a high-confidence protein interaction network including a predicted interaction between the dengue NS5 protein and mTOR (20). To validate and further study this interaction, we performed co-immunoprecipitation experiments using exogenously expressed NS5. GFP-tagged NS5 or GFP alone was transfected into 293FT cells, leading to modest expression of the fusion protein in the cytosol with nuclear accumulation, as has been previously reported (21). Protein complexes were then affinity purified using GFP-specific nanobodies (22). Subsequent western blotting of affinity purified NS5 complexes revealed mTOR, Raptor, and Rictor proteins, demonstrating that NS5 interacts with both mTORC1 and mTORC2 (Fig. 1).

**FIGURE 1.**
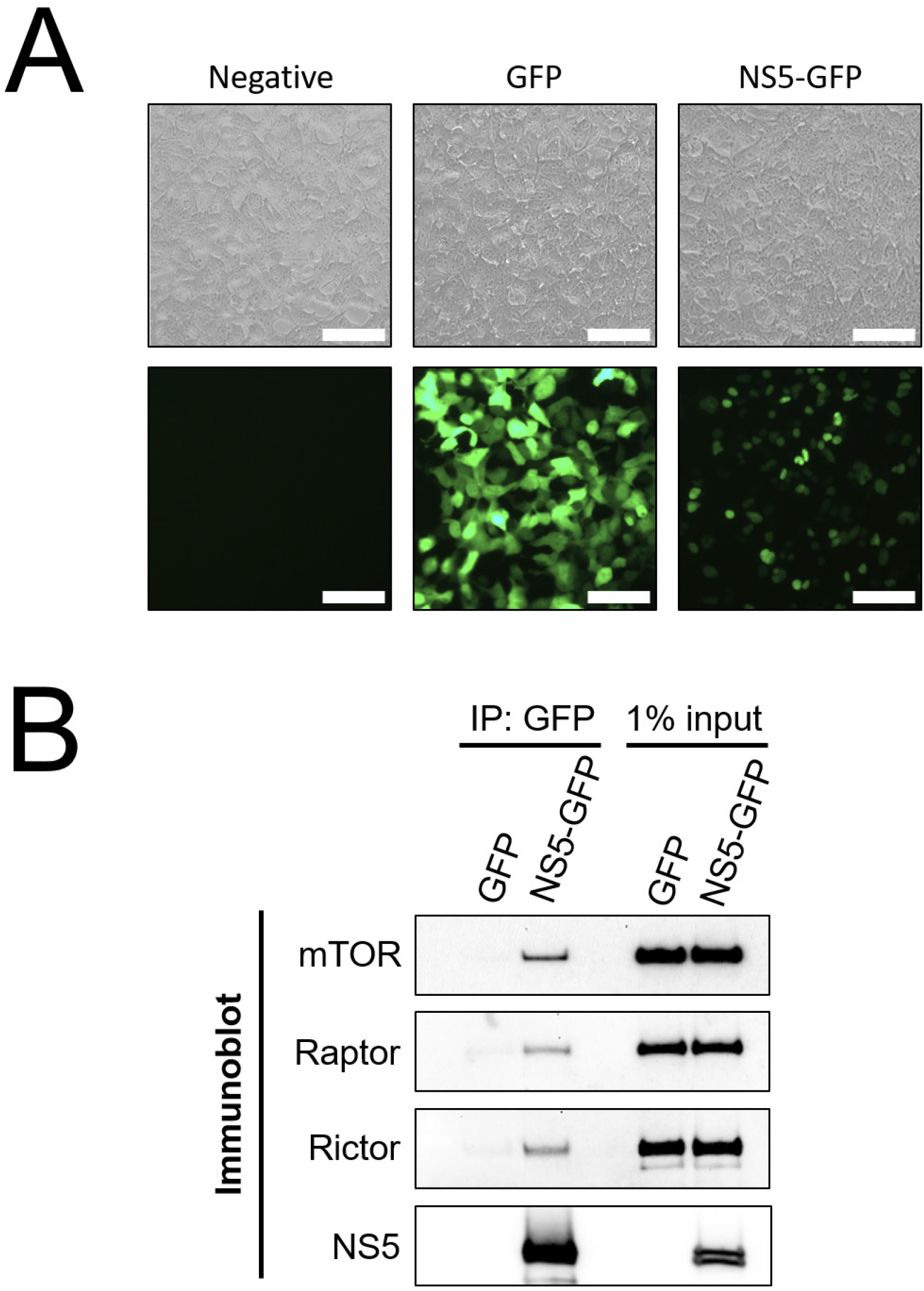
Dengue NS5 interacts with mTORC1 and mTORC2. *A*, Expression of NS5-GFP fusion protein. 293FT cells were transfected with expression plasmids encoding GFP or NS5-GFP fusion protein, and live cells were imaged by fluorescence microscopy. Bar = 100 μm. *B*, Cells were transfected with GFP or NS5-GFP plasmids as in (*A*), and lysates were affinity captured using anti-GFP nanobodies. 50% of the eluates or 1% of the input lysate were then analyzed by western blot analysis with the indicated antibodies.

### mTORC2 is activated during dengue replication and is required for efficient viral replication

To investigate what impact NS5 interactions may have on dengue viral infection, and to explore the differential effect of NS5 interaction with mTORC1 and mTORC2, the expression of each complex was silenced using lentivirus-expressed shRNA targeting mTOR protein, Raptor (a component of mTORC1) or Rictor (a component of mTORC2) or, as a control, a nonspecific scrambled oligo-sequence (23). We used HepG2 hepatoma cells for these experiments as they have been used extensively in studies of dengue replication, apoptosis, autophagy and ER stress (15, 20, 24, 25). shRNA transduction resulted in substantially decreased protein abundance of mTOR, Raptor and Rictor (Fig. 2*A*). To assess whether Raptor and Rictor knockdown inhibited the signaling activity of mTORC1 and mTORC2, we examined the phosphorylation status of well-characterized downstream targets (p70 S6K thr389 for mTORC1 and AKT ser473 for mTORC2) (23, 26). Knockdown of Raptor led to diminished mTORC1 signaling as evidenced by decreased S6K thr389 phosphorylation, and knockdown of Rictor led to diminished mTORC2 signaling as evidenced by decreased AKT ser473 phosphorylation (Fig. 2*A*). We observed that depletion of mTORC1 led to reciprocal activation of mTORC2 activity, consistent with prior reports suggesting that mTORC1 represses mTORC2 signaling (27) (Fig. 2*A*). The converse did not appear to be the case, as depletion of mTORC2 did not lead to increased phosphorylation of S6K by mTORC1 (Fig. 2*A*). Furthermore, mTOR knockdown diminished levels of Raptor and Rictor, but knockdown of neither Raptor or Rictor influenced levels of the other (Fig. 2*A*).

**FIGURE 2.**
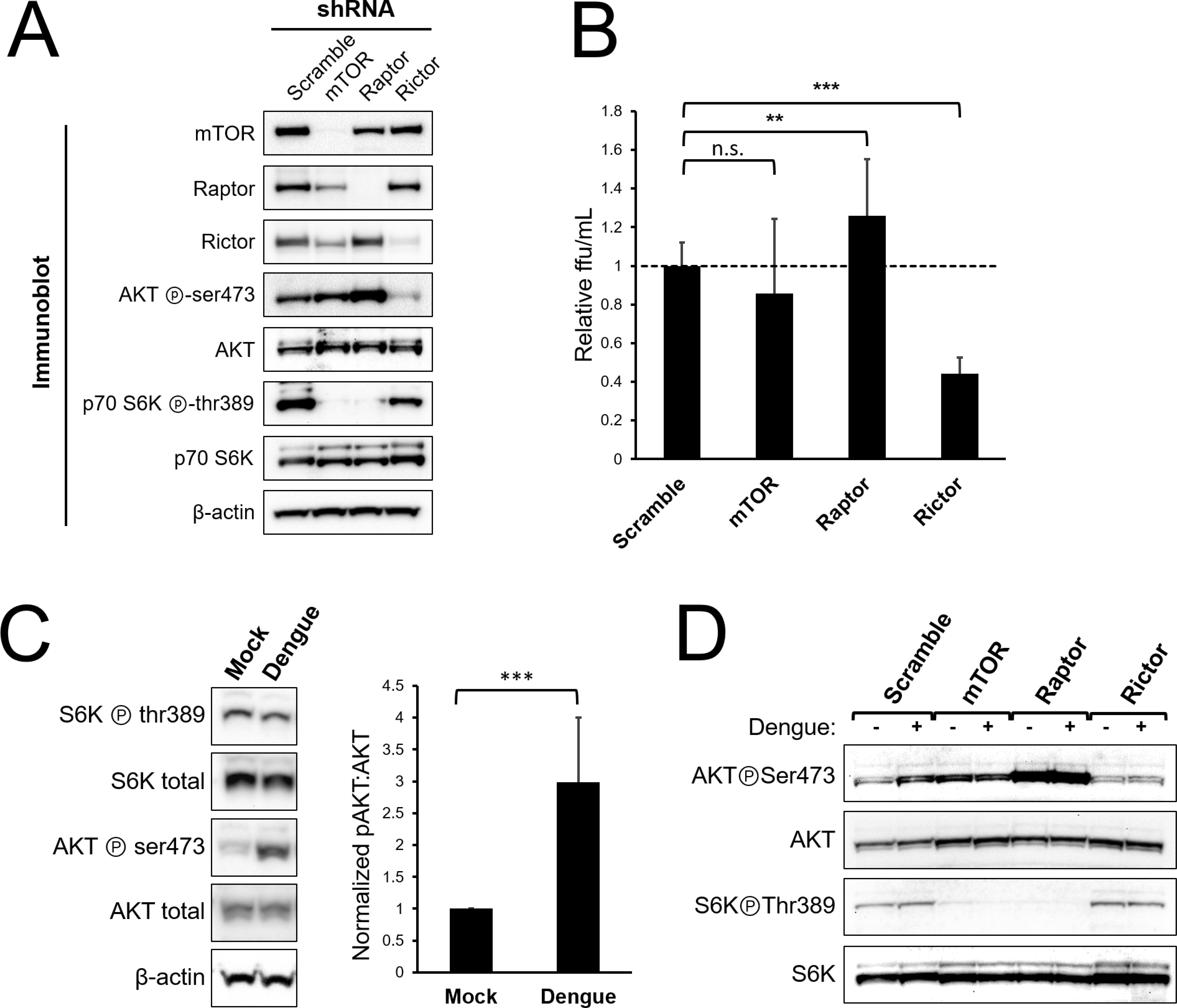
Dengue infection activates mTORC2 signaling and is required for maximal viral replication. *A*, Selective inactivation of mTORC1 and mTORC2 in HepG2 cells. Cells were transduced with lentivirus encoding shRNA directed against mTOR protein, Raptor, or Rictor, or with a non-specific scramble sequence, and then selected with puromycin. Lysates were analyzed by western blot with the indicated antibodies. *B*, mTORC2 inactivation diminishes dengue replication. Cells were transduced with lentiviral shRNA vectors as in (*A*) and were then infected with dengue MON601 at a MOI of 1. 24 h later cell culture supernatants were collected and titrated on Vero cells. The data represent scramble-normalized values from 4 independent experiments. *C*, mTORC2 is activated during dengue infection. HepG2 cells were infected with dengue at MOI of 4. Cells were collected at 36 h post-infection and analyzed by western blot with the indicated antibodies. The bar graph shows the ratio of phospho-AKT to total AKT band intensity averaged from 7 independent experiments. *D*, Rictor knockdown abrogates mTORC2 activation by dengue. Cells were transduced with shRNA encoding lentivirus as in (*A*) and were infected with dengue at MOI of 4. Cells were collected at 36 hpi and were analyzed by western blot with the indicated antibodies. Error bars are one standard deviation. p values are derived from 2-tailed Student’s t test; n.s. denotes p > 0.05, * denotes p < 0.05, ** denotes p < 0.01, and *** denotes p < 0.001.

We next assessed the effect of reduced mTOR protein, Raptor (mTORC1), and Rictor (mTORC2) on dengue serotype 2 replication by infecting respective knockdown cells and quantifying the amount of infectious virus released. Interestingly, Rictor knockdown led to a substantial decrease in the amount of viral replication, while Raptor knockdown led to a small but significant increase in replication. No significant effect on viral replication was observed when mTOR protein was knocked down (Fig. 2*B*).

Having observed a decrease in viral replication with mTORC2 inactivation, we next asked whether dengue infection affected the activity of mTORC2. To do this, we infected HepG2 cells with dengue and assessed the phosphorylation status of the downstream targets AKT and S6K by western blot. In these experiments, mTORC2-specific phosphorylation at AKT ser473 was increased in infected cells compared to mock treated cells, and the ratio of ser473 phosphorylated AKT to total AKT was consistently increased in infected cells (Fig. 2*C*, p < 0.001). Using mTOR, Raptor and Rictor knockdown cells, we again observed an increase in AKT ser473 phosphorylation in infected cells, which was abrogated in Rictor knockdown cells, demonstrating that mTORC2 is required for dengue-induced AKT phosphorylation. As in Fig. 2*A*, knockdown of mTOR protein or Raptor led to increased mTORC2 activity, which was not further increased by dengue infection (Fig. 2*D*).

### mTORC2 inhibition does not affect cellular morphology, growth rates or dengue-induced LC3-II accumulation

The increase in mTORC2 activity during dengue infection and the requirement for mTORC2 activity for maximal viral replication led us to investigate potential physiologic functions of mTORC2 that are exploited by the virus. Because mTORC2 has been shown to play a role in regulating cell proliferation (28), we considered the possibility that our findings could be explained by altered cell growth rates. We assessed cell growth rates in knockdown cells by measuring cell counts and CFSE dilution. No significant differences in growth rates were observed upon comparison of mTORC2 knockdown cells with control cells (Fig. 3*A* and *B*). Because mTOR has also been shown to play a role in the maintenance of cell morphology by modulating ion channel activity and altering dynamics of the actin cytoskeleton (3, 4), we examined the morphology of the mTORC2-depleted cells and their actin cytoskeleton. Cells treated with shRNA targeting Rictor showed normal cellular morphology, normal cell spreading and a morphologically normal actin cytoskeleton (Fig. 3*C*), suggesting that mTORC2 inactivation was not causing derangements in cell architecture in this setting.

**FIGURE 3.**
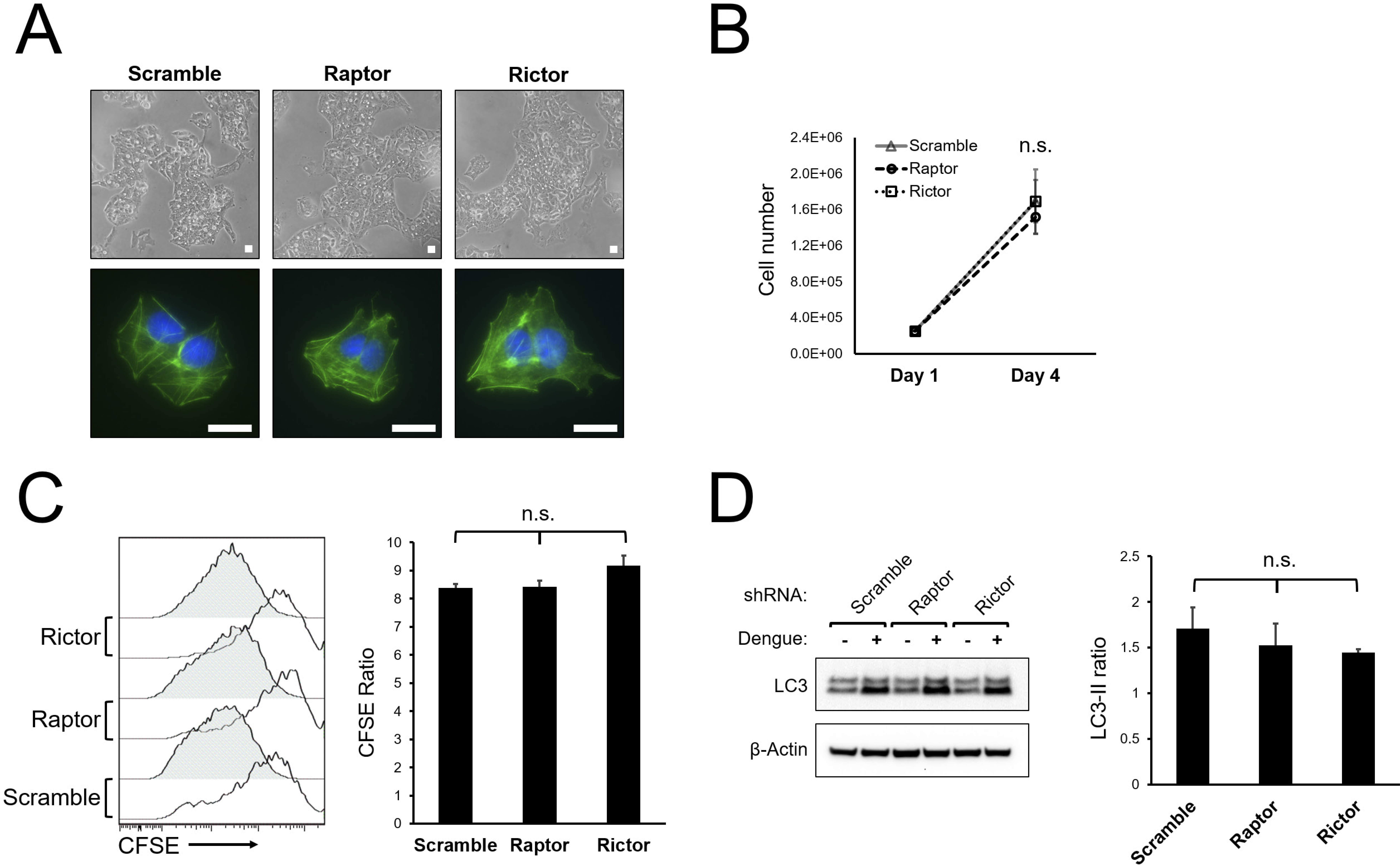
mTORC2 inactivation does not alter HepG2 cell morphology, cytoskeletal architecture, growth rate, or LC3 lipidation in response to dengue infection. *A*, HepG2 cells were transduced with lentiviral vectors encoding shRNA directed against mTOR protein, Raptor or Rictor, or with a non-specific scramble sequence and were selected with puromycin. Cells were imaged by light microscopy (top panels) or stained with fluorophore-labeled phalloidin and imaged with fluorescence microscopy (bottom panels). Bar = 100 μm *B*, Cells were transduced with shRNA lentiviral vectors as in (*A*) and were plated at equal densities. Cells were then counted using an automated cell counter 4 days later. Data represent cell counts from 4 separate cultures. *C*, Cells were transduced with shRNA lentiviral vectors as in (*A*) and were plated at equal densities. Cells were then loaded with CFSE and analyzed by flow cytometry 24 h (open histograms) and 96 h (shaded histograms) later. Bar graph represents the ratio of CFSE mean fluorescence intensity between the 24 and 96 h timepoint, averaged from 4 separate cultures. *D*, mTORC2 inhibition does not alter dengue-mediate LC3-II lipidation. Cells were transduced with shRNA lentiviral vectors as in (*A*) and were infected with dengue at MOI 4. Cells were harvested at 36 h postinfection and lysates were analyzed by western blot with the indicated antibodies. The bar graph shows the ratio of LC3-II band intensity between dengue infected and mock treated cells from 3 independent experiments. Error bars are one standard deviation. p values are derived from 2-tailed Student’s t test; n.s. denotes p > 0.05, * denotes p < 0.05, ** denotes p < 0.01, and *** denotes p < 0.001.

Although autophagy is canonically regulated by mTORC1, we also considered the possibility that inactivation of mTORC2 suppresses dengue replication by altering autophagy through indirect regulatory effects on mTORC1. While the effect that dengue has on the movement of cargo through the canonical autophagy pathway is unclear, it is well known that dengue induces the accumulation of autophagosomes, characterized by accumulation of lipidated LC3 protein (LC3-II) (10, 12, 13, 15). To assess effects of mTORC2 inhibition on the pro-autophagic activity of dengue, we measured LC3-II isoform levels in dengue-infected knockdown cells and observed similar degrees of dengue-induced LC3-II accumulation (Fig. 3*D*), suggesting that mTORC2 inactivation does not block the effects of dengue on autophagy.

### mTORC2 inactivation causes increased dengue-mediated apoptosis and cell death

We next asked whether mTORC2 could function as a pro-survival factor in infected cells, given the known role for mTORC2 in regulating cell survival (28–30). We infected Rictor knockdown or control cells with dengue for 24 or 36 h and, as a positive control for apoptosis, treated the cells with staurosporine. We then asked whether apoptosis or cell death was altered in the context of mTORC2 inhibition by measuring activated caspase 3 expression and cell viability within infected and uninfected cells using flow cytometry. At 24 hours post-infection (hpi), we observed a similar frequency of infected cells when comparing control cells to Rictor knockdown cells, arguing against an early block in viral replication in mTORC2-deficient cells. However, at 36 hpi, Rictor knockdown was associated with a significant decrease in the proportion of infected cells (Fig. 4*A* and *B*). Assessment of activated caspase 3 expression and cell viability at 24 h revealed a small increase in apoptosis and cell death in mTORC2-deficient cells, which were not statistically significant (Fig. 4*A*, *C*, and *D*). However, by 36 hpi, a marked increase in apoptosis and cell death was observed in infected mTORC2 knockdown cells, both of which were highly significant (4*A*, *C*, and *D*, p < 0.001 for activated caspase 3 level, p < 0.01 for cell viability). Interestingly, while staurosporine treatment increased cell death and apoptosis at similar levels in both scramble and Rictor knockdown cells, suggesting that the increased apoptosis seen in dengue-infected cells was specific for virus-induced apoptosis. We also measured the release of dengue from infected cells and found that less infectious virus was released from Rictor knockdown cells, and that the difference increased from 24 to 36 hpi, indicating that virus production is diminished as apoptotic cell death increases.

**FIGURE 4.**
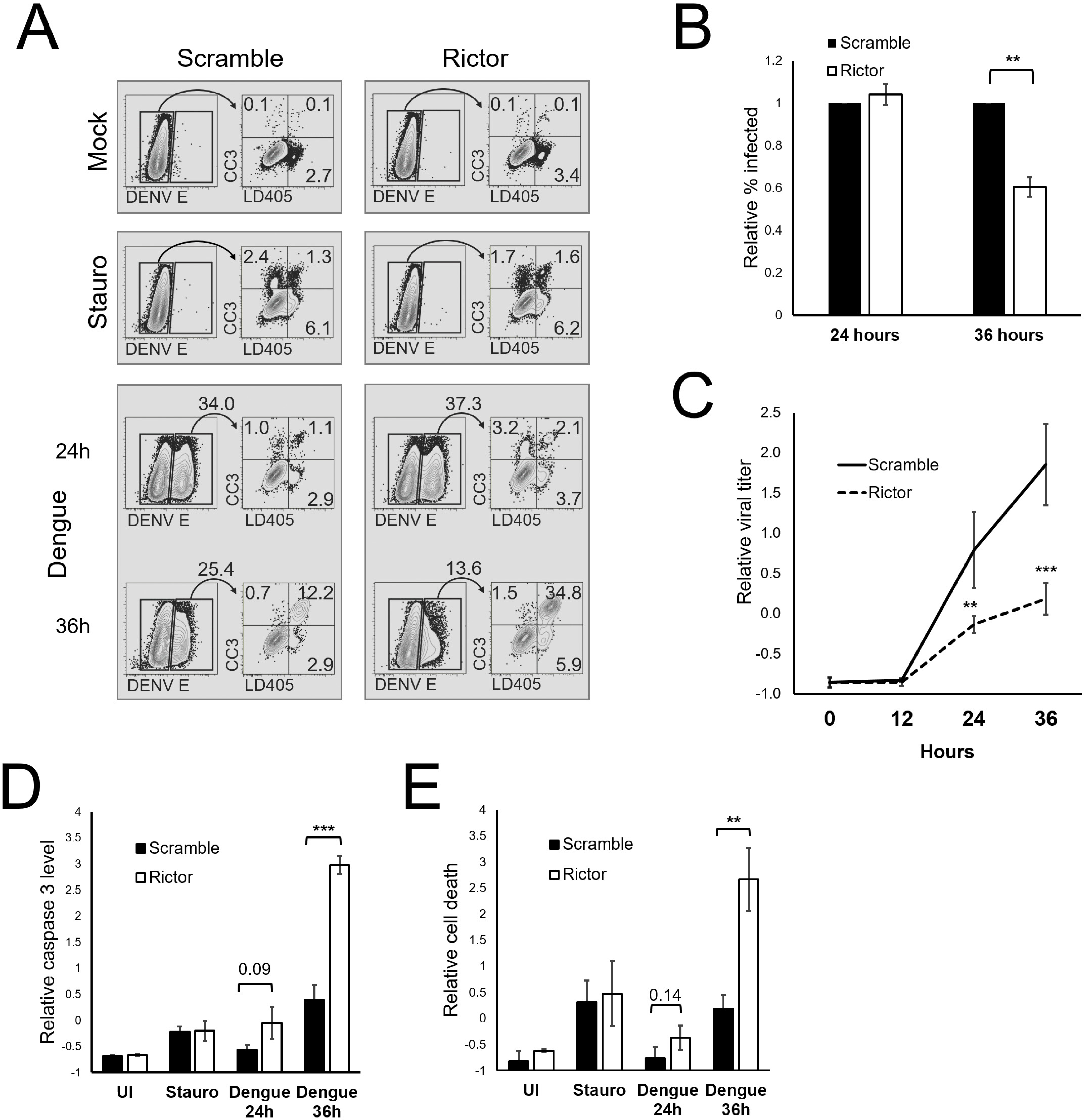
mTORC2 inhibition leads to increased dengue-induced apoptosis and cell death. *A*, HepG2 cells were infected with dengue at MOI of 4 for 24 or 36 h, mock treated, or treated with 5 μM staurosporine overnight. Cells were then collected and stained with a cell-impermeable amine reactive dye (LD405), followed by fixation, permeabilization, and staining with anti-activated caspase 3 and anti-flavivirus E protein antibodies. Infected (E-positive) cells were gated and activated caspase 3 (CC3) and LD405 expression were analyzed within that population. Numbers represent population frequencies in each gate/quadrant. *B*, Cells were treated as in (*A*), and the percent of infected (E-positive) cells was calculated. The bar graph shows the average values for 3 independent experiments, each normalized to the scramble condition. *C*, Cells were treated as in (*A*), and the percent of infected cells expressing activated caspase 3 was calculated. The bar graph represents Z-normalized average values from 3 independent experiments. *D*, Cells were treated as in (*A*), and the percent of infected cells positive for LD405 was calculated. The bar graph represents *Z*-normalized values from 3 independent experiments. p values are derived from 2-tailed Student’s t test; n.s. denotes p > 0.05, * denotes p < 0.05, ** denotes p < 0.01, and *** denotes p < 0.001.

## DISCUSSION

In this study, we describe a role for host mTORC2 in dengue replication, demonstrating that dengue can interact with the cellular mTOR signaling system, activating mTORC2 and thus promoting cell survival and permitting efficient viral replication. Specifically, the data we present here suggest that mTORC2 plays a critical role in supporting viral production by counteracting virus-induced apoptosis of the host cell. The induction of apoptosis in dengue infection has been well-described, occurring in several cell types both *in vitro* and *in vivo*, including endothelial cells, dendritic cells, and hepatocytes (31–34). The finding of apoptotic cells in human autopsy specimens from severe dengue cases has led to speculation that apoptosis contributes to pathogenesis in these cases (32). While apoptosis may contribute to tissue damage and pathogenesis from the host perspective, it is also an important mechanism for host control of viral infection, triggering cell death before infectious progeny can be released (35) and shaping the subsequent immune response. Since uncontrolled apoptosis would be detrimental to viral replication, it is not surprising that dengue has evolved a mechanism to attenuate the induction of apoptosis in infected cells. In the present study, we found that decreasing mTORC2 signaling in infected cells leads to an increased frequency of apoptosis in cell death, and the induction of apoptosis corresponds to a decrease in the release of viral progeny from infected cells. Interestingly, the susceptibility of mTORC2-deficient cells to apoptosis was specific for dengue infection in our experiments, as neither baseline apoptosis nor apoptosis in response to staurosporine treatment increased over control upon depletion of mTORC2. Furthermore, we found that dengue infection triggers the activation of mTORC2, which likely represents a viral adaptation to maintain cell survival during infection. While this strategy has not been described in other viral infections, it has been reported as mechanism for cancer cell survival and metastasis in several malignancies (36).

Dengue NS5 protein binds to mTOR, suggesting that the viral protein modulates mTOR signaling during infection. NS5 appears to bind both mTORC1 and mTORC2, evidenced by the co-immunoprecipitation of Raptor and Rictor with NS5. The molecular consequences of NS5-mTOR interaction remain to be investigated. One possibility is that NS5 binding to mTORC2 directly facilitates activation of the complex or acts as an adaptor protein to stabilize interactions with downstream targets of the complex. Given that there is crosstalk between mTORC1 and mTORC2 signaling (37), it is also possible that interaction of NS5 with mTORC1 could activate mTORC2, either via de-repression of mTORC2 or through alterations in mTOR protein stoichiometry.

The findings we present here also highlight the role of proteomic approaches in understanding virus-host interactions. Much of the recent focus in dengue research has utilized high-throughput genetic screens employing approaches such as RNAi and CRISPR (38–40). While these genetic screens have identified host dependency factors, because they rely on measuring viral gene expression or cell survival as the readout for infection resistance, they are likely biased toward host factors involved in the early stages of infection such as viral entry and gene expression as opposed to host factors needed for late infection events such as viral release, maturation, and host cell survival. For this reason, it is perhaps not surprising that components of the autophagy machinery and mTOR signaling have not been consistently identified in these screens, despite the roles of these host factors reported here and by others (14, 15). Moreover, genetic screens stop short of identifying specific molecular interactions between virus and host that mediate regulatory processes, which can be elucidated when interactions between viral and host proteins are interrogated via proteomics. Nonetheless, it is notable that Rictor inhibition was associated with decreased relative infection in 2 RNAi libraries previously reported, although that finding did not meet the authors’ criteria for further validation (40). In contrast to genetic screens, proteomics methods can identify host interactions at all stage of the viral life cycle, and subsequent validation can identify important host interactions. When combined with quantitative approaches to distinguish high-probability interactors from non-specific binding, proteomics approaches can identify host factors with high validity (20).

The observation that mTORC2 is required for efficient dengue replication raises the possibility of mTOR as a target for host-directed antiviral therapeutic development. mTOR is a highly “druggable” target, and intense interest in the mTOR pathways in oncology and neuropathology fields has spurred the development of multiple new small molecule inhibitors with high specificity for mTOR (29, 41). In the solid organ transplant infectious disease field, data have begun to emerge that certain mTOR inhibitors (used as anti-organ rejection medications) may have anti-viral properties, reducing the risk of some viral reactivation syndromes (42). In the case of dengue *perse*, candidate compounds would likely need to have mTORC2 specificity, since mTORC1 inhibition appears to enhance viral replication. While most newer generation mTOR inhibitors have dual mTORC1/C2 specificity, a recently developed compound CID613034 has been described that specifically inhibits mTORC2, demonstrating the feasibility of specific mTORC2 inhibition (43). Since multiple biochemical steps occur between mTORC2 assembly and the resultant anti-apoptotic outcome, components of the mTORC2 signaling cascade might also provide useful targets for host-based interventions.

The role of mTORC2 signaling in the pathogenesis of other viral infections remains to be determined; however, there is evidence that other viruses including West Nile, influenza and human cytomegalovirus may also stimulate mTORC2 signaling (16–18). Whether mTORC2 is important for viral replication and serves as an anti-apoptotic mechanism in those viruses remains to be determined, but it is possible that mTORC2 modulation is a common mechanism used by several viruses to counteract the host’s programmed cell death response. If that is the case, host-directed therapeutic interventions targeting mTORC2 could be active against multiple viral pathogens.

## MATERIALS AND METHODS

### Antibodies and reagents

The following antibodies were obtained from Cell Signaling Technology (Danvers, MA) and used at 1:1000 for western blotting: Raptor clone 24C10, Rictor clone D16H9, phospho-Rictor thr1135 clone D30A6, mTOR 7C10, AKT clone 11E7, phospho-AKT ser473 clone D9E (used at 1:2000), p70 S6K clone 49D7, phospho-p70 S6K thr389 clone 108D2, LC3A/B clone D3U4C, and B-actin clone 13E5. The dengue 2 E protein antibody PA5-32246 was used at 1:10,000 for western blotting (Thermo-Fisher Scientific, Waltham, MA). Secondary detection for western blotting used anti-rabbit HRP antibody diluted 1:10,000 (Amersham ECL, GE Healthcare, Chicago, IL). Polyclonal affinity purified NS3 and NS5 proteins were produced as previously described (20). The pan-flavivirus E antibody 4G2 was prepared from hybridoma supernatants and purified by protein A/G chromatography. For flow cytometry experiments, 4G2 was conjugated to FITC according to the manufacturer’s instructions (FluoroTag kit, Sigma-Aldrich, St. Louis, MO). The dimerized GFP nanobody construct LaG-16–G4S–LaG-2 (green lobster) was a gift from Michael Rout, and was prepared as previously described (22). Phalloidin-Alexafluor 488 was obtained from Thermo-Fisher scientific and used according to their instructions. Staurosporine was obtained from Millipore Sigma (Burlington MA) and was used at 5 μM concentration.

### Plasmids

The lentiviral shRNA plasmids pLKO.1 scramble, Raptor_2, mTOR_2, and Rictor_2 were gifts from David Sabatini (Addgene plasmids 1864, 1858, 1856, and 1854). pMD2.G and psPAX2 were gifts from Didier Trono (Addgene plasmids 12259 and 12260). Dengue NS5-GFP fusion construct was generated by inserting the dengue New Guinea C NS5 coding sequence from pDVW601 (44) into pACGFP1-N1 (Clontech) as previously described (20).

### Cell culture

HepG2 and Vero cells were cultured at 37 °C in 5% CO_2_, in medium composed of MEM supplemented with 10% FBS, 1× non-essential amino acids, 50 units/mL of penicillin and 50 μg/mL of streptomycin. Medium was replenished frequently during experiments to avoid signaling changes caused by nutrient or growth factor depletion. The *Aedes albopictus* derived cell line C6/36 was propagated at 28 °C in 5% CO_2_ in medium composed of MEM supplemented with 10% FBS, 1× non-essential amino acids, 50 units/mL of penicillin and 50 μg/mL of streptomycin. The B lymphocyte hybridoma cell line D1-4G2-4-15 was obtained from the ATCC (Manassas, VA), and was maintained in ATCC Hybri-Care Medium supplemented with 10% FBS and 1.5 g/L sodium bicarbonate.

### Transfections

293FT cells were plated to 70-80% density in antibiotic free medium. Transfection complexes were prepared by mixing plasmid DNA with polyethyleneimine (PEI Max 40k, Polysciences Inc., Warrington, PA) in a 1:4 mass ratio. After 15 min incubation at room temperature DNA complexes were added dropwise to the cell cultures.

### Affinity capture

To prepare GFP affinity-capture beads, the dimerized GFP nanobody LaG-16–G4S–LaG-2 (green lobster) was covalently linked to magnetic beads (Dynabeads M-270 epoxy Thermo Fisher Scientific) as previously described (45). Briefly, 5 μg of nanobody were used per 1 mg of Dynabeads, with conjugations carried out in 0.1 M sodium phosphate buffer and 1 M ammonium sulfate, with an 18-to 20-h incubation at 30 °C. Beads were then washed sequentially with 0.1 M sodium phosphate buffer, 100 mM glycine pH 2.5, 10 mM Tris-HCl pH 8.8, 100 mM triethylamine, 1× PBS (4 times), PBS + 0.5% Triton X-100, and 1× PBS. For affinity capture experiments, cells were harvested 48 h after transfection. Cells were washed with ice cold PBS and then lysed with 1% Triton X-100, 0.5% sodium deoxycholate, 110 mM potassium acetate pH 7.5, 20 mM HEPES, 2 mM MgCl_2_, 25 mM NaCl, and 1× protease/phosphatase inhibitor cocktail (Cell Signaling Technology). Lysates were clarified by centrifugation for 10 min at 13,000 × *g* at 4 °C. Affinity capture beads were immediately added to the clarified lysate and incubated for 10 min at room temperature with rotation. Beads were then washed 3× with lysis buffer, and bound protein complexes eluted with 1.1× LDS sample buffer for 10 min at 70 °C. For SDS-PAGE and western blot analysis, 10× reducing agent (Thermo-Fisher Scientific) was added and samples were heated for an additional 10 min at 70 °C. SDS-PAGE and western blot analysis were performed as described below. Gel staining was performed using Sypro Ruby fluorescent gel stain (Thermo-Fisher Scientific) according to the manufacturer’s instructions.

### shRNA-mediated gene silencing

To generate lentiviral vector stocks, shRNA constructs were cotransfected with pMD2.G and psPAX2 into HEK 293T cells. Supernatants were harvested, passed through 0.45 μm filters, layered on 20% sucrose cushions, and centrifuged at 100,000 × *g* for 4 h at 4 °C. Lentiviral pellets were resuspended in OptiMEM and stored at −80 °C until use. For lentiviral transductions, viral stocks were diluted to the desired concentration with OptiMEM and 0.8 μg/mL polybrene and added to cells. At 48 h post-transduction, cells containing stably integrated constructs were selected using 2 μg/mL puromycin. Experiments were performed on cell lines that were maintained and passaged for no more than 3 weeks before discarding and establishing fresh cell lines.

### Virus and infections

DENV-2 MON601, a molecular clone of DENV-2 New Guinea strain C[46], was generated by transfection of *in vitro*-transcribed RNA into Vero cells, followed by no more than 5 passages in C6/36 cells. Virus was propagated by infecting 80% confluent C6/36 monolayers with low-passage stock virus at a MOI of 0.01, and harvesting infectious supernatants 5-7 days post-infection. Infectious supernatants were cleared of cellular debris by centrifugation and stored at −80 °C until use. Virus stocks and experimental infectious supernatants were titrated using a flow cytometry approach which has been described elsewhere (46). Briefly, serially diluted virus stocks were used to infected Vero cells in a multi-well plate. Cells were harvested 20-24 h post-infection, fixed and permeabilized, and stained with 4G2-FITC. The percentage of infected cells was then used to calculate the number of fluorescence forming units (FFU) per milliliter of inoculum. For experimental infections, virus was diluted in OptiMEM to the desired MOI and incubated on cells for 90 min at 37 °C. Virus was then removed, the cells washed, and complete growth medium added.

### Western blot analysis

Cells were placed on ice and washed with ice-cold PBS. Cells were then collected and lysed with Triton X-100 lysis buffer (1% Triton X-100, 120 mM NaCl, 1 mM EDTA, 40 mM HEPES pH 7.4, and 1× protease and phosphatase inhibitor cocktail (Cell Signaling Technology)). To prepare whole cell extracts, SDS lysis buffer (2% SDS, 50 mM Tris pH 7.4, 5% glycerol, 5 mM EDTA, 1 mM NaF, 1 mM DTT, and 1× phosphatase & protease inhibitor cocktail) was used instead of Triton X-100 lysis buffer. Protein concentration was determined using BCA assay and a BSA standard curve, and equivalent amounts of protein were mixed with 4× LDS sample buffer and 10× reducing agent (Thermo-Fisher Scientific), followed by denaturation at 70 °C for 10 min. Proteins were then resolved on 4-12% Bis-Tris gels (for lower molecular weight proteins) or 3-8% Tris-Acetate gels (for higher molecular weight proteins) (NuPage, Thermo-Fisher Scientific) and run in MOPS or Tris-Acetate running buffer respectively. Proteins were transferred to PVDF membrane and blocked in 5% milk/TBS-T for 1-2 h. Primary antibodies were diluted in 5% BSA/TBS-T, and incubated overnight at 4 °C. Membranes were washed and incubated with anti-rabbit HRP antibody for 1-2 h at room temperature. Membranes were then washed with TBS-T, exposed to chemiluminescent substrate, and imaged using a digital CCD platform (Fluorchem E, Protein Simple, San Jose, CA). Band densitometry was performed using ImageJ software.

### Fluorescence microscopy

Cells were fixed with 4% paraformaldehyde/PBS for 15 min at room temperature, permeabilized with 0.1% Triton X-100/PBS for 10 min at room temperature and blocked with 5% normal goat serum in 0.05% Tween-20/PBS. Cells were then stained with phalloidin-Alexafluor 488 (Thermo Fisher Scientific) for 1 h at room temperature and mounted using medium containing DAPI.

### Flow cytometry

For viability analysis, cells were trypsinized, washed with PBS, and stained with a cell impermeable amine reactive dye (LIVE/DEAD Violet A.K.A. LD405, ThermoFisher Scientific) according to the manufacturer’s instructions. Cells were fixed and permeabilized using the BD Cytofix/Cytoperm kit according to the manufacturer’s instructions (BD Biosciences). Permeabilized cells were stained with fluorophore-conjugated antibodies as indicated in the text. Cells were analyzed on a BD LSR-II cytometer, and data were analyzed using FlowJo software.

## ACKNOWLEDGMENTS

F.D.M was supported by a postdoctoral fellowship of the Canadian Institutes for Health Research. This study was supported by grant R01GM101183 to A.K. and grants R21AI124266 and P41GM109824 to J.D.A. from the National Institutes of Health.

